# CAZymes in *Maribacter dokdonensis* 62-1 from the Patagonian shelf: Genomics and physiology compared to related flavobacteria and a co-occurring *Alteromonas* strain

**DOI:** 10.1101/2020.12.08.416198

**Authors:** Laura A. Wolter, Maximilian Mitulla, Jovan Kalem, Rolf Daniel, Meinhard Simon, Matthias Wietz

## Abstract

Carbohydrate-active enzymes (CAZymes) are an important feature of bacteria in productive marine systems such as continental shelves, where phytoplankton and macroalgae produce diverse polysaccharides. We herein describe *Maribacter dokdonensis* 62-1, a novel strain of this flavobacterial species, isolated from alginate-supplemented seawater collected at the Patagonian continental shelf. *M. dokdonensis* 62-1 harbors a diverse array of CAZymes in multiple polysaccharide utilization loci (PUL). Two PUL encoding polysaccharide lyases from families 6, 7, 12 and 17 allow substantial growth with alginate as sole carbon source, with simultaneous utilization of mannuronate and guluronate as demonstrated by HPLC. Furthermore, strain 62-1 harbors a mixed-feature PUL encoding both ulvan- and fucoidan-targeting CAZymes. Core-genome phylogeny and pangenome analysis revealed variable occurrence of these PUL in related *Maribacter* and *Zobellia* strains, indicating specialization to certain “polysaccharide niches”. Furthermore, lineage- and strain-specific genomic signatures for exopolysaccharide synthesis possibly mediate distinct strategies for surface attachment and host interaction. The wide detection of CAZyme homologs in algae-derived metagenomes suggests global occurrence in algal holobionts, supported by sharing multiple adaptive features with the hydrolytic model flavobacterium *Zobellia galactanivorans*. Comparison with *Alteromonas* sp. 76-1 isolated from the same seawater sample revealed that these co-occurring strains target similar polysaccharides but with different genomic repertoires, coincident with differing growth behavior on alginate that might mediate ecological specialization. Altogether, our study contributes to the perception of *Maribacter* as versatile flavobacterial polysaccharide degrader, with implications for biogeochemical cycles, niche specialization and bacteria-algae interactions in the oceans.

## INTRODUCTION

Continental shelves are productive marine systems, where photosynthesis by pelagic phytoplankton and benthic macroalgae yields considerable amounts of organic matter. Polysaccharides constitute a major fraction of the algae-derived organic matter, with important roles in nutrient cycles and microbial metabolism (Hehemann et al. 2014, Arnosti et al. 2021). Consequently, diverse bacteria are specialized for the degradation of algal polysaccharides, colonization of algal surfaces and other types of biological interactions (van der Loos et al. 2019, Wolter et al. 2020, Ferrer-González et al. 2020).

Cultured bacterial strains are a valuable resource for studying the ecological and biogeochemical implications of microbial polysaccharide degradation, complementing molecular and metagenomic approaches on community level (Arnosti et al. 2011, Wietz et al. 2015, Matos et al. 2016, Reintjes et al. 2017, Grieb et al. 2020). Culture-based studies revealed the diversity and functionality of carbohydrate-active enzymes (CAZymes), which encompass polysaccharide lyases (PL), glycoside hydrolases (GH), carbohydrate-binding modules (CBM), carbohydrate esterases (CE), glycosyl transferases (GT) and auxiliary carbohydrate-active oxidoreductases (Lombard et al. 2014). CAZyme-encoding genes are frequently transferred between microbes, providing effective mechanisms of adaptation and niche specialization (Hehemann et al. 2016).

Flavobacteria, including the families *Flavobacteriaceae and Cryomorphaceae*, are major players in marine polysaccharide degradation. Comparable to the human gut, marine flavobacteria can degrade various polysaccharides through dedicated genetic machineries (Teeling et al. 2012, Fernández-Gómez et al. 2013). Flavobacterial CAZymes are typically clustered with *susCD* genes in polysaccharide utilization loci (PUL) for orchestrated uptake and degradation (Grondin et al. 2017). For instance, the marine flavobacterium *Zobellia galactanivorans* has complex biochemical and regulatory mechanisms for degrading laminarin, alginate, agar and carrageenan (Hehemann et al. 2012, Thomas et al. 2012, 2017, Labourel et al. 2014, Ficko-Blean et al. 2017, Zhu et al. 2017). Comparable abilities have been described in the flavobacterial genera *Formosa agarivorans* and *Gramella forsetii* through diverse PUL (Mann et al. 2013, Kabisch et al. 2014, Reisky et al. 2019). Also *Maribacter*, the “sister genus” of *Zobellia*, exhibits hydrolytic activity (Bakunina et al. 2012, Zhan et al. 2017). Accordingly, both *Maribacter* and *Zobellia* are abundant on macroalgal surfaces (Martin et al. 2015), and related PUL have been detected during phytoplankton blooms in the North Sea (Kappelmann et al. 2019). Furthermore, several *Maribacter* and *Zobellia* strains stimulate algal development by producing morphogenesis factors (Matsuo et al. 2005, Weiss et al. 2017). Hence, both genera are important from ecological and biotechnological perspectives.

Here, we describe CAZyme content and hydrolytic capacities of *Maribacter dokdonensis* strain 62-1, isolated from an alginate-supplemented microcosm at the Patagonian continental shelf (Wietz et al. 2015). This highly productive marine region harbors frequent phytoplankton blooms and abundant coastal macroalgae, indicating regular availability of polysaccharides (Acha et al. 2004, Garcia et al. 2008). Our CAZyme characterization in the pangenomic context illustrates the role of CAZymes, PUL and exopolysaccharide-related genes in niche specialization among *Maribacter* and *Zobellia*. The finding of diverse algae-adaptive traits, together with detection of CAZyme homologs in macroalgae-derived metagenomes, highlight *Maribacter* spp. as a versatile associate of marine algae. Notably, *Maribacter dokdonensis* 62-1 has been isolated from the same sample as *Alteromonas* sp. 76-1 with shown capacity for alginate and ulvan degradation (Koch et al. 2019b), illustrating that distantly related hydrolytic strains co-occur in the same habitat. Comparison of their CAZyme machineries illuminated whether these strains might employ different ecophysiological strategies or compete for resources. These eco-evolutionary perspectives into CAZyme diversity and corresponding niche specialization contribute to the understanding of ecophysiological adaptations behind community-level polysaccharide degradation (Teeling et al. 2012, 2016). Considering the abundance and biogeochemical relevance of algal polysaccharides, our study adds further evidence to the eco-evolutionary role of CAZyme in marine flavobacteria.

## MATERIALS AND METHODS

### Isolation and cultivation

Strain 62-1 was isolated in April 2012 from a microcosm with surface seawater collected at the Patagonian continental shelf (47° 56’41”S 61° 55’23”W) amended with 0.001% sodium alginate (Wietz et al., 2015). Purity was confirmed by PCR amplification of the 16S rRNA gene after several rounds of subculturing. Alginate utilization was analyzed in seawater minimal medium (SWM) (Zech et al. 2009) supplemented with 0.2% sodium alginate (cat. no. A2158; Sigma-Aldrich, St. Louis, MO) as sole carbon source in comparison to SWM + 0.4% glucose. Precultures were grown from single colonies for 24h, washed three times with sterile SWM, and adjusted to an optical density of 0.1 measured at 600nm (OD600). Main cultures were inoculated with 1% (v/v) of washed preculture in triplicate, followed by cultivation at 20°C and 100 rpm with regular photometric measurements (diluted if OD600 >0.4).

### Substrate quantification

At each OD measurement, subsamples of 5 mL were filtered through 0.22 μm polycarbonate filters into combusted glass vials and stored at ⎼20°C. Alginate concentrations were quantified by High Performance Liquid Chromatography (HPLC) of its monomers mannuronate and guluronate after chemical hydrolysis (20h, 100°C, 0.1 M HCl) in combusted and sealed glass ampoules. Samples were neutralized with 6 N NaOH, desalted using DionexOnGuard II Ag/H cartridges (Thermo Fisher Scientific, Waltham, MA), and eluted with 100 mM sodium acetate tri-hydrate in 100 mM NaOH. Concentrations were determined in three dilutions per sample (0.01, 0.002, 0.001%) using a Carbopac PA 1 column (Thermo Fisher Scientific) and pulsed amperometric detection according to (Mopper et al. 1992). A calibration curve was generated using hydrolyzed 1% alginate solution (*R*^2^ = 0.97). Glucose concentrations were measured using samples diluted to 0.001% with MilliQ followed by HPLC with NaOH (18 mM) as eluent and a Carbopac PA 1 column (Thermo Fisher Scientific). A calibration curve was generated using 24 concentrations from 0.025−10 μM glucose (*R*^2^ = 0.99).

### Genome sequencing and taxonomy

Genomic DNA was extracted using the PeqGold DNA Isolation Kit (PEQLAB, Germany) according to the manufacturer’s instructions. The genome was sequenced with Illumina technology using a GAIIx sequencing platform on paired-end libraries prepared with the Nextera XT DNA Kit (Illumina, San Diego, CA). A total of 84 contigs (0.5 to 330 kb, average 55 kb) were assembled using SPAdes v3.0 (Bankevich et al. 2012), followed by error correction using BayesHammer (Nikolenko et al. 2013), gene prediction using the IMG pipeline (Markowitz et al. 2012) and conversion to EMBL format using EMBLmyGFF3 (Norling et al. 2018). The draft genome is available at ENA (https://www.ebi.ac.uk/ena) and IMG (https://img.jgi.doe.gov) under PRJEB40942 and 2588253514, respectively. In addition, the genome has been deposited under https://github.com/matthiaswietz/Maribacter. Phylogenetic analysis was carried out with 92 core genes identified using UBCG (Na et al. 2018), with *Capnocytophaga ochracea* DSM 7271 as outgroup. The resulting nucleotide alignment was visually confirmed for consistency and the best substitution model (GTR+G) computed using ModelTest-NG (Darriba et al. 2020). A maximum-likelihood phylogeny with 1000 bootstrap replicates was calculated using RaxML v8.2 (Stamatakis 2014) on the CIPRES Science Gateway (Miller et al. 2010).

### Comparative genomics

Genomes of 62-1 and related strains (Supplementary Table S1) were compared using bioinformatic software. Average nucleotide identities were calculated using the Enveomics web application (Rodriguez-R and Konstantinidis, 2016). Core, accessory and unique genes were identified from protein-translated genes using OrthoFinder (Emms & Kelly 2019) using a 30% identity cutoff. CAZymes were identified using dbCAN2 (Zhang et al., 2018), only considering hits with e-value <10^−15^ and >65% query coverage. Gene annotations and PUL boundaries were manually curated based on the CAZy and UniprotKB-Swissprot databases (Lombard et al. 2014, Bateman et al. 2017). Sulfatases were identified using SulfAtlas v1.1 (Barbeyron et al. 2016a), only considering hits with e-value <10^−1^ and >40% query coverage. Genes were assigned to KEGG classes and pathways using KAAS and KEGG Mapper (Moriya et al. 2007, Kanehisa & Sato 2020). PUL homologies were analyzed by custom-BLAST in Geneious Pro v7 (https://www.geneious.com) and PULDB (Terrapon et al. 2018). Genes for downstream processing of alginate monomers (*kdgA*, *kdgF*, *kgdK* and *dehR*) were identified by searching homologs from *Gramella forsetii* (NCBI assembly GCA_000060345.1). Putative *kduI* and *kduD* genes for processing unsaturated uronates were identified by searching homologs from *Gramella flava* (NCBI assembly GCA_001951155.1). Signal peptides and CRISPR-Cas modules were predicted using SignalP v5.0 (Almagro Armenteros et al. 2019) and CRISPRCasFinder (Couvin et al. 2018) respectively.

### Statistical evaluation and data visualization

Data were processed in RStudio (https://rstudio.com) using R v3.6 (R Core Team 2018) and visualized using packages ggplot2, PNWColors and pheatmap (Wickham 2016, Kolde 2018, Lawlor 2020). Code and files for reproducing the analysis are available under https://github.com/matthiaswietz/Maribacter.

## RESULTS AND DISCUSSION

Strain 62-1 was isolated from alginate-supplemented seawater collected at the Patagonian continental shelf (Wietz et al., 2015). Colonies on solid medium are round, smooth and yellowish-colored. Genome sequencing resulted in a draft genome (84 contigs) with a cumulative length of 4.6 Mb, encoding 4,105 predicted proteins. Core genome-based phylogeny revealed clear assignment to *Maribacter* from the *Flavobacteriaceae* (Fig. 1), with 99.8% 16S rRNA gene similarity and 97.8% average nucleotide identity to *Maribacter dokdonensis* DSW-8^T^ (Supplementary Fig. S1). Hence, strain 62-1 is a novel member of this flavobacterial species. *M. dokdonensis* DSW-8^T^ originates from South Korean and hence subtropical waters, demonstrating occurrence of closely related strains on global scales.

**Fig. 1.**
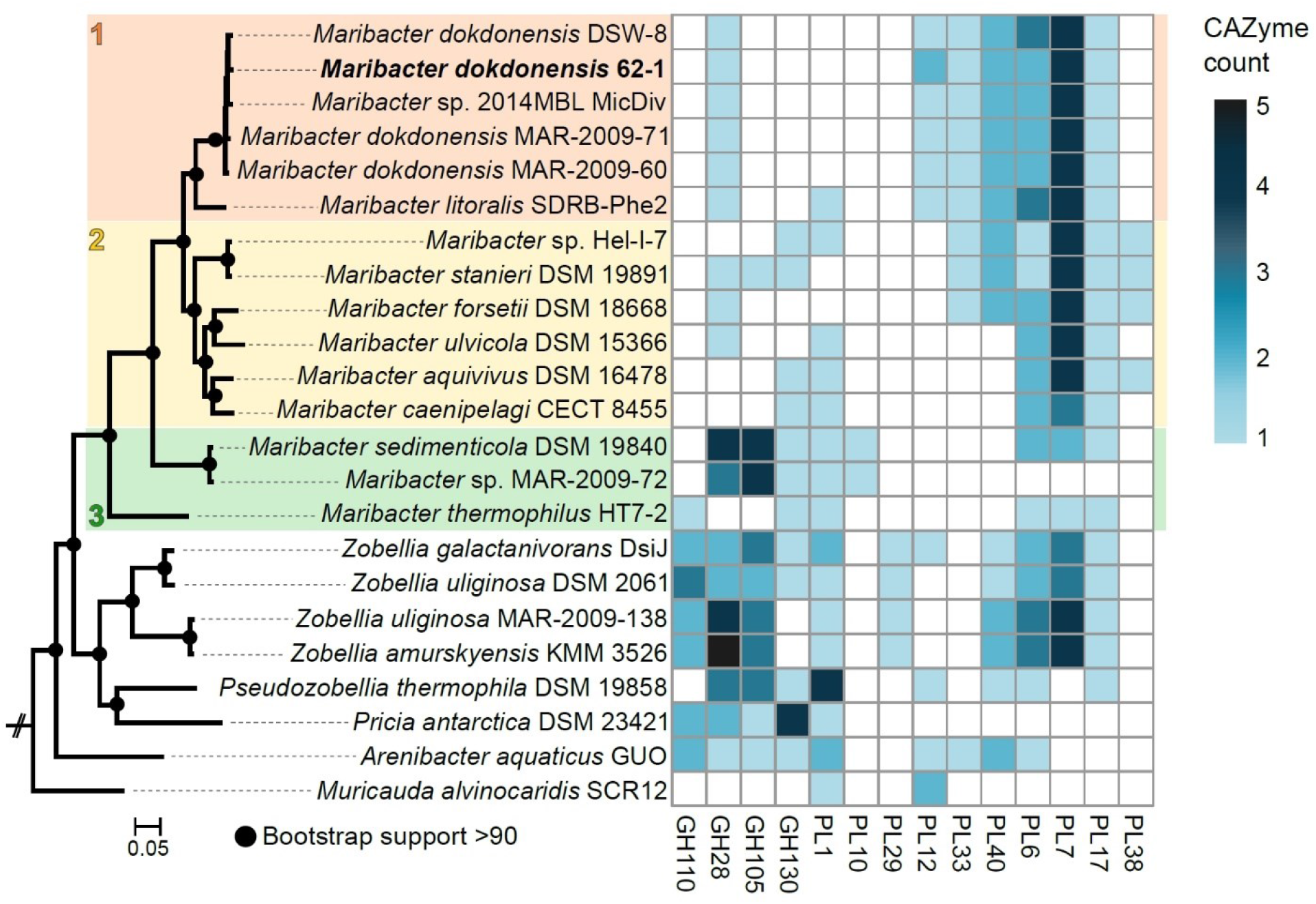
Maximum-likelihood phylogeny based on 82 core genes (left panel) and numbers of genes from selected CAZyme families (right panel) in *Maribacter dokdonensis* 62-1 and related strains. *Capnocytophaga ochracea* DSM 7271 served as outgroup. The three resolved *Maribacter* lineages are numbered and colored. PL: polysaccharide lyase; GH: glycoside hydrolase.

### CAZymes in the phylogenomic context

*Maribacter dokdonensis* strain 62-1 encodes 90 CAZymes, corresponding to 2% of all protein-encoding genes (Table 1, Supplementary Table S1). As described in detail below, CAZymes commonly clustered with *susCD* genes, the hallmark of PUL in Bacteroidetes (Grondin et al. 2017). The presence of twelve polysaccharide lyases from PL families 6, 7, 12 and 17 illustrates specialization towards alginate, confirmed by physiological experiments (Fig. 2). Most PL12 are classified as heparinases, but co-localization with known alginate lyase families indicates alginolytic activity. Strain 62-1 furthermore encodes PL33 and PL40 lyases that potentially target ulvan (Table 1). CAZyme numbers and diversity match the hydrolytic potential of related *Maribacter* and *Zobellia* strains included for comparison (Supplementary Table S1), corroborating the adaptation of these taxa to algal substrates and surfaces (Bakunina et al. 2012, Martin et al. 2015, Kwak et al. 2017, Zhan et al. 2017, Chernysheva et al. 2019).

**Table 1.** CAZyme families encoded by *Maribacter dokdonensis* 62-1. PL: polysaccharide lyase; GH: glycoside hydrolase; GT: glycosyl transferase; CBM: carbohydrate-binding module; CE: carbohydrate esterase; AA: auxiliary carbohydrate-active oxidoreductase.

| CAZyme class | CAZyme family |
| --- | --- |
| PL | 6, 7, 12, 17, 33, 40 |
| GH | 1, 2, 3, 5, 10, 13, 15, 16, 20, 23, 28, 29, 31, 32, 43, 65, 73, 78, 85, 88, 95, 97, 109, 113, 117, 141, 144 |
| GT | 2, 4, 5, 9, 19, 20, 26, 27, 28, 30, 51, 56, 83 |
| CBM | 9, 48, 67 |
| CE | 1, 11, 14 |
| AA | 1 |

**Fig. 2.**
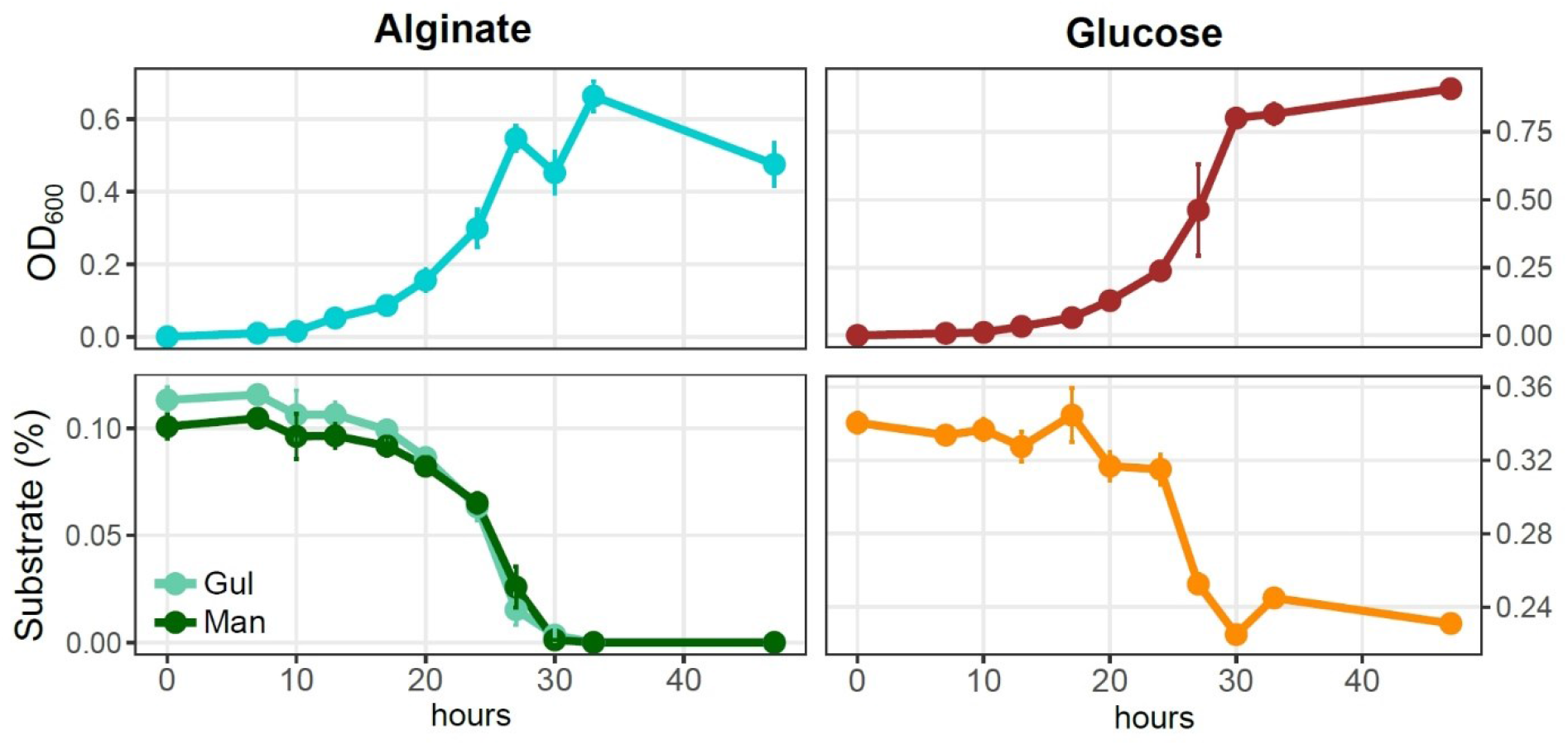
Growth of *Maribacter dokdonensis* 62-1 with alginate compared to glucose as sole carbon source, illustrated by optical density (upper panels) and substrate utilization as determined by HPLC (lower panels). Gul: guluronate; Man: mannuronate.

We contextualized CAZyme patterns with phylogenetic relationships (Fig. 1, Supplementary Table S2) and a general overview of the *Maribacter* pangenome (Supplementary Table S3). Core genome-based phylogeny resolved three lineages, each with distinct signatures of CAZymes and exopolysaccharide-related genes (Fig. 1). *Maribacter* lineages 1 and 2 encode more PLs than lineage 3 (Wilcoxon rank-sum test, *p* = 0.01), including an entire additional PUL for alginate degradation. Moreover, one PL7 and one PL12 are unique to lineage 1 (see detailed paragraph below). In contrast, a sizeable PUL encoding a PL10 pectate lyase with CE8 methylesterase domain is restricted to lineage 3 (Fig. 1). This PL10-CE8 presumably corresponds to a specific pectin-related niche, supported by elevated numbers of GH families 28 and 105 that participate in pectinolytic activity (Hobbs et al. 2019).

Notably, lineages 2 and 3 harbor distinct clusters for the biosynthesis of cell wall exopolysaccharides (EPS). Furthermore, specific O-antigen and GT variants occur on sublineage- and strain-level (Supplementary Table S3), presumably facilitating surface colonization and specific interactions with algal hosts (Deo et al. 2019). This marked variability on fine phylogenetic levels suggests EPS-related genes as important adaptive feature among *Maribacter* strains, mediating different surface-attachment and host-interaction mechanisms (Lee et al. 2016, Decho & Gutierrez 2017).

### Alginate degradation relates to two alginolytic PUL

The array of alginate lyases in *Maribacter dokdonensis* 62-1 allowed considerable growth with alginate as sole nutrient source (Fig. 2). Growth was only slightly lower than with the monosaccharide glucose, signifying an excellent adaptation for polysaccharide degradation. HPLC demonstrated that both monomeric building blocks of alginate, mannuronate (M) and guluronate (G), were utilized at the same rate (Fig. 2). Hence, strain 62-1 efficiently degrades the complete alginate polymer without preferring one of its monomers.

Alginolytic abilities correspond to genes in two major PUL plus additional, single alginate lyase genes dispersed throughout the genome (Fig. 3, Supplementary Table S2). AlgPUL1 harbors four adjacent PL6−12−6−17 lyase genes, co-localized with a *susCD* pair and all genes for downstream processing of alginate monomers (Fig. 3A). Hence, AlgPUL1 presumably encodes the complete metabolic cascade from external polymer breakdown (PL6, PL12), oligosaccharide hydrolysis (PL17) to monomer processing (*kdgA*, *kdgF*, *kdgK* and *dehR*). The co-localization of PL families with structural and catalytic diversity (Xu et al. 2017) presumably facilitates access to various alginate architectures, e.g. relating to polymer length or the ratio between M and G in poly-G, poly-M or mixed stretches. Upstream of AlgPUL1 is a cluster encoding one GH3 and two GH144 genes, together with *susCD* and sulfatase genes (Fig. 3). The two regions are separated by a type II-C CRISPR-Cas locus, signifying a mobile genomic region that might facilitate the exchange of adjacent CAZymes. Both GH144 genes share ~50% amino acid identity to an endo-glucanase of *Chitinophaga pinensis* (Abe et al. 2017), indicating glucosidase or endoglucanase activity. Their occurrence in all *Maribacter* and *Zobellia* (orthologous groups 0002512 and 0002513; Supplementary Table S3), many marine flavobacteria as determined by BLASTp (data not shown) as well as human gut microbes (McNulty et al. 2013) indicates ecological relevance across diverse habitats. The smaller AlgPUL2 encodes two PL7 lyases (Fig. 3B) and might be auxiliary to AlgPUL1, potentially being activated by particular M/G architectures or environmental signals (Lim et al. 2011). The prediction of different signal peptide variant in the two PL7 (Supplementary Table S2) indicated that one PL7 is freely secreted whereas the other is anchored to the cell membrane. This complementary extracellular localization might boost alginolytic activity.

**Fig. 3:**
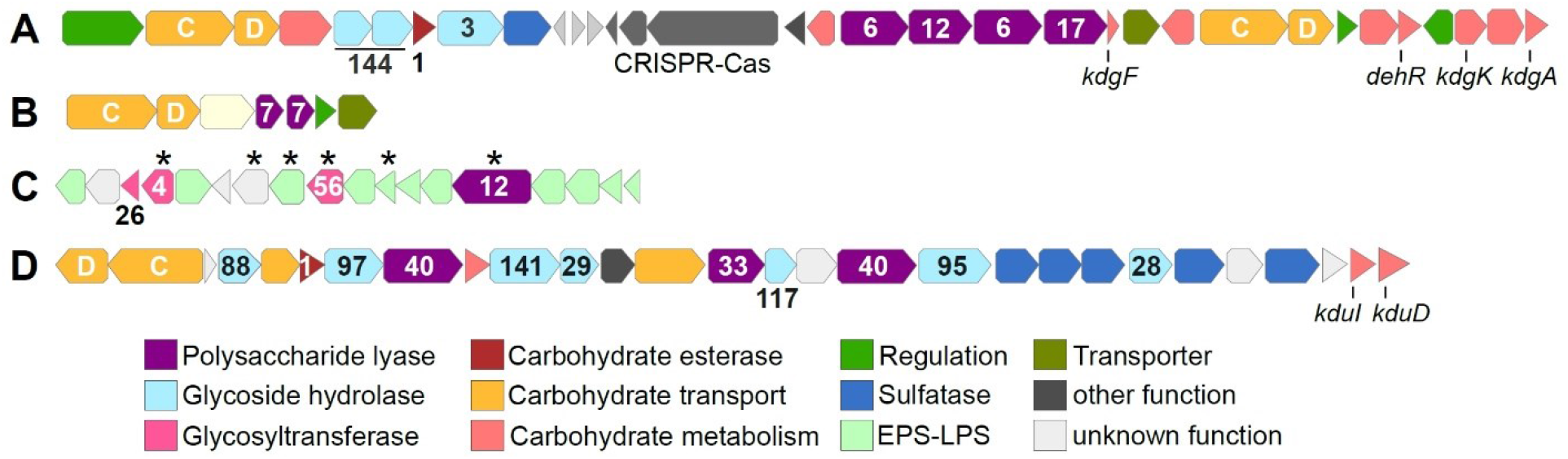
Major PUL in *Maribacter dokdonensis* 62-1. Numbers designate CAZyme families, letters *susCD* genes, and asterisks unique genes of strain 62-1. *kdgA*: 2-keto-3-deoxy-D-gluconate aldolase; *kdgF*: gene for 2-keto-3-deoxy-D-gluconate linearization; *kdgK:* 2-keto-3-deoxy-D-gluconate kinase; *dehR*: 4-deoxy-L-erythro-5-hexoseulose uronate reductase; *kduI*: 5-keto 4-deoxyuronate isomerase; *kduD*: 2-deoxy-D-gluconate 3-dehydrogenase.

Comparative genomics revealed homologs of AlgPUL1 in most related *Maribacter* and *Zobellia* strains, whereas AlgPUL2 is restricted to *Maribacter* lineages 1 and 2. The observed structural diversity of alginolytic PUL among *Maribacter* and *Zobellia* genomes overall corresponds to core-genome relationships, supporting the notion of lineage-specific genomic signatures (Supplementary Fig. S2). Notably, a PL12 only occurs in the AlgPUL1 variant of lineage 1 (Fig. 1). This PL was presumably acquired from distant Bacteroidetes taxa, considering 65% amino acid identity to FNH22_29785 from *Fulvivirga* sp. M361 (Cytophagales). In contrast, lineages 2 and 3 encode a PL7 in this position, or solely harbor the PL6 and PL17 (Supplementary Fig. S2). An almost identical AlgPUL2 occurs in *Flavivirga eckloniae* (locus tags C1H87_08155−08120), with 65% amino acid identity between the respective PL7 lyases. The original isolation of *F. eckloniae* from the alginate-rich macroalga *Ecklonia* (Lee et al. 2017) highlights this PUL as adaptation to algae-related niches. In *Zobellia*, AlgPUL1 and AlgPUL2 occur in mixed combinations and in distant genomic regions (Thomas et al. 2012), suggesting internal recombination events (Supplementary Fig. S2).

*Maribacter dokdonensis* 62-1 encodes a unique PL12 (locus tag 00457) within an exopolysaccharide-related cluster, including several unique GTs (Fig. 3C). This cluster might represent a link between the degradation and biosynthesis of polysaccharides, considering that other bacteria regulate EPS production and release via PLs or GHs (Bakkevig et al. 2005, Köseoğlu et al. 2015). The PL12 has 39% amino acid identity to ATE92_1054 of the North Sea isolate *Ulvibacter* sp. MAR-2010-11 (Kappelmann et al. 2019), indicating ecological relevance of horizontally transferred PL12 homologs in distant habitats. The PL12 is among the two lyases without predicted signal peptide (Supplementary Table S2) and hence likely retained within the cell, supporting an intracellular role in EPS metabolism.

### A PUL related to ulvan and fucoidan

Strain 62-1 harbors additional PUL for the degradation of other algal polysaccharides. The co-localization of two PL40 lyases, one PL33 from subfamily 2 as well as several sulfatases indicates targeting a sulfated polysaccharide, presumably ulvan (Fig. 3D). Although PL33 are classified as chondroitin or gellan lyases, our findings suggest an extended substrate range, potentially specific to subfamily 2. The two PL40 variants (locus tags 01347 and 01356) only have 30% identity and hence do not originate from duplication. Notably, PL40_01347 lacks a signal peptide compared to PL40_01356 and the PL33 (Supplementary Table S2), suggesting different secretory behavior and complementary functionality. The PL40_01347 homolog is conserved in most *Maribacter* and *Zobellia* genomes (Supplementary Fig. S2), whereas PL40_01356 and the PL33 are missing in *Zobellia*. The latter PLs share 70% and 77% amino acid identity with a CAZyme pair in *Formosa agariphila* (locus tags BN863_10330 and BN863_10340 respectively), originally annotated as heparinase and PL8 but confirmed as PL33 and PL40 using the latest CAZy database (Mann et al. 2013, Lombard et al. 2014). Their absence in *Zobellia* but presence in distantly related *Arenibacter* spp. (Fig. 1) indicates separate acquisition after speciation, potentially via transfer events between *Formosa* and *Maribacter*.

The combination of PL33 and PL40 differs from ulvanolytic PUL in other bacteria, which largely comprise ulvan lyases from PL families 24, 25 and 28 (Foran et al. 2017, Reisky et al. 2018, Koch et al. 2019b). Furthermore, the PUL in strain 62-1 also encodes α-fucosidases, including GH29, GH95, GH97, GH117 and GH141 comparable to a CAZyme plasmid in the verrucomicrobium *Lentimonas* (Sichert et al. 2020). Furthermore, a co-localized GH28 might remove galacturonic acid from fucoidan (Sichert et al. 2020), further metabolized by adjacent *kduI* and *kduD* genes into the Entner-Doudoroff pathway (Salinas & French 2017). These observations indicate an alternative hydrolytic activity towards fucoidan, supported by fucose importer, fuconolactonase and fuconate dehydratase genes (locus tags 01371, 01373, 01376 respectively) encoded downstream of the PUL.

### Ecological implications

To establish a broader ecological context, we searched homologs of the PL33-PL40 pair (specific to *Maribacter*) and the PL12 within the EPS gene cluster (unique to strain 62-1) in 102 microbial metagenomes from marine plants. The co-detection of PL33 and PL40 homologs on diverse algae and seagrasses from global locations (Fig. 4A; Supplementary Table S4) illustrates that relatives of strain 62-1 occur in related niches worldwide. The presence of PL33-PL40 homologs on brown, green and red macroalgae indicates that predicted hydrolytic capacities are functional *in situ* and support establishment in algal holobionts. The brown macroalgae *Macrocystis* and *Ecklonia* are rich in alginate whereas ulvan can constitute ~40% of the green algae *Ulva* (Kidgell et al. 2019), highlighting the importance of related CAZymes for associated bacteria and why *Maribacter* spp. are common algal epibionts (Martin et al. 2015).

**Fig. 4:**
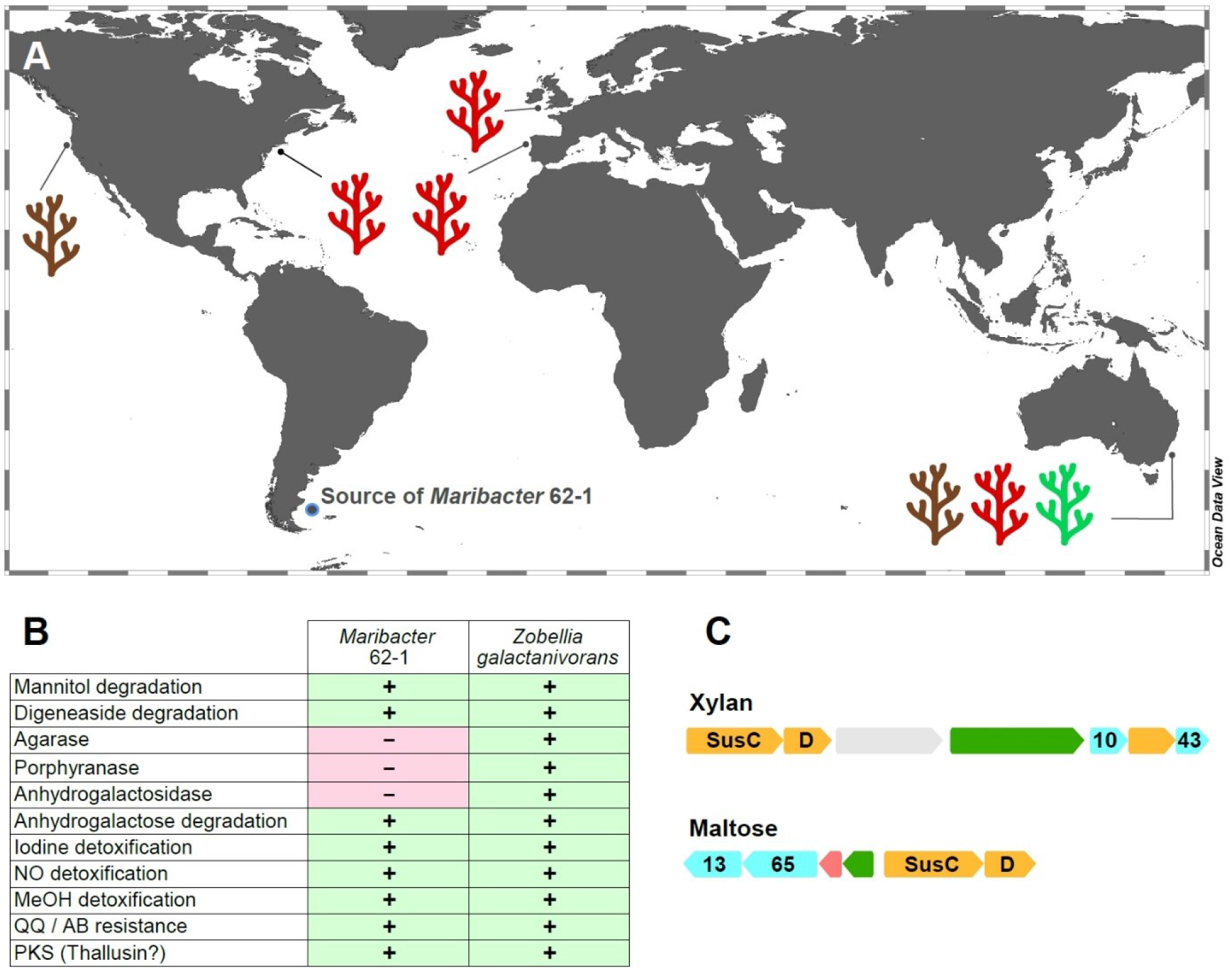
Ecological implications of CAZyme diversity in *Maribacter dokdonensis* 62-1. **A:** Detection of polysaccharide lyase homologs in macroalgae-derived metagenomes from global locations. Colors indicate brown, red or green algal taxa (see Supplementary Table S4 for details). **B:** Presence (+) or absence (−) of algae-adaptive traits compared to characterized features in *Zobellia galactanivorans*. MeOH: methanol, QQ: quorum quenching; AB: antibiotic; PKS: polyketide synthase. **C:** Gene clusters encoding glycoside hydrolases (cyan), *susCD* (orange), regulators (green) and carbohydrate-processing genes (red) with homology to PUL in *Z. galactanivorans* targeting xylan and maltose (locus tags 0158−0164 and 1248−1253 respectively). Numbers designate GH families.

Wide detection of the PL12 unique to strain 62-1 (Supplementary Table S4) supports the presumed role of the lyase and adjacent EPS genes in surface attachment. This gene arrangement might permit the formation of specific biofilm structures, helping to reduce competition with co-existing bacteria. EPS on algal surfaces could also provide a protective matrix, minimizing diffusion of secreted CAZymes and retaining hydrolysis products for maximal uptake (Vetter et al. 1998). In turn, EPS could also be advantageous in pelagic waters from which 62-1 has been isolated, where aggregation on self-produced EPS could constitute a protective refugium and facilitate survival when algal substrates are unavailable (Decho & Gutierrez 2017). In general, the diversity of EPS genes on fine phylogenetic levels might mediate distinct interactions with algal hosts, considering that EPS can be strain-specific determinants of host interaction (Lee et al. 2016, Deo et al. 2019).

In addition to metagenomic analyses, we searched strain 62-1 for genes encoding characterized algae-adaptive traits in *Zobellia galactanivorans*, a closely related hydrolytic flavobacterium (Barbeyron et al. 2016b). This approach identified homologous genes for degradation of mannitol and digeneaside in strain 62-1 (Fig. 4B; Supplementary Table S4). Furthermore, we found GH43-GH10 and GH13-GH65 gene pairs targeting the algal carbohydrates xylan and maltose (Fig. 4C). We also detected most homologs for anhydrogalactose utilization and hence the second step in carrageenan metabolism, but no GH127 or GH129 anhydrogalactosidase genes for initial hydrolysis (Ficko-Blean et al. 2017). Hence, *Maribacter* might utilize hydrolysis products from primary degraders, employing a secondary “harvester” strategy for carrageenan on red macroalgae (Hehemann et al. 2016). However, agarase or porphyranase homologs were not detected (Fig. 4B), illustrating an overall narrower niche range than *Zobellia*.

In addition to carbohydrate utilization, we found other features typical for interactions with macroalgae. For instance, genes for the detoxification of iodine and nitrous oxide (Fig. 4B; Supplementary Table S4) likely counteract algal defense mechanisms (Ogawa et al. 1995, Verhaeghe et al. 2008). Detoxification of superoxide and hydrogen peroxide is associated with two dismutases and 21 thioredoxin-like oxidoreductases (Supplementary Table S4), an almost two-fold higher count despite an ~1Mb smaller genome than *Zobellia* (Barbeyron et al. 2016b). The predicted ability to detoxify methanol is likely advantageous during demethylation of pectinous algal substrates (Koch et al. 2019a). Moreover, strain 62-1 encodes traits that might be advantageous to algal hosts. For instance, a putative homoserine lactone lyase (Fig. 4B; Supplementary Table S4) might interfere with communication of bacterial competitors and prevent their biofilm formation. This process, often termed quorum quenching, might antagonize resource competitors and modulate the composition of the holobiont (Wahl et al. 2012). Presence of a PKS might allow biosynthesis of thallusin, an essential algal morphogen produced by both *Zobellia* and *Maribacter* (Matsuo et al. 2003, Weiss et al. 2017). Another conserved feature is the complete pathway for biosynthesis of biotin (vitamin B7; locus tags 03353‒03359 in strain 62-1), a common characteristic of macroalgal epibionts (Karimi et al. 2020). Strain 62-1 misses a single gene in biosynthetic pathways for riboflavin (vitamin B2) and pantothenate (B5), but might provide intermediates to other bacteria encoding the complete pathway (Paerl et al. 2018, Shelton et al. 2019).

### Comparison with *Alteromonas* sp. 76-1 from the same habitat

The gammaproteobacterial strain *Alteromonas* sp. 76-1, isolated from the same seawater sample as *Maribacter dokdonensis* 62-1, has a comparable predisposition towards polysaccharide degradation (Koch et al. 2019b). The presence of alginolytic and ulvanolytic systems in both 62-1 and 76-1 supports the notion that alginate and ulvan commonly occur at the Patagonian continental shelf. Although the two strains might hence compete for these resources, their corresponding CAZyme repertoires substantially differ. While sharing similar numbers of PL6 and PL7 alginate lyase genes (Fig. 5A), PL12 are restricted to *Maribacter* and PL18 to *Alteromonas*, indicating different alginolytic strategies. Furthermore, concerning ulvanolytic activity, *Alteromonas* encodes PL24 and PL25 lyases on a CAZyme plasmid opposed to PL33 and PL40 on the *Maribacter* chromosome (Fig. 5A), potentially influencing the regulation and transfer of these genes.

**Fig. 5.**
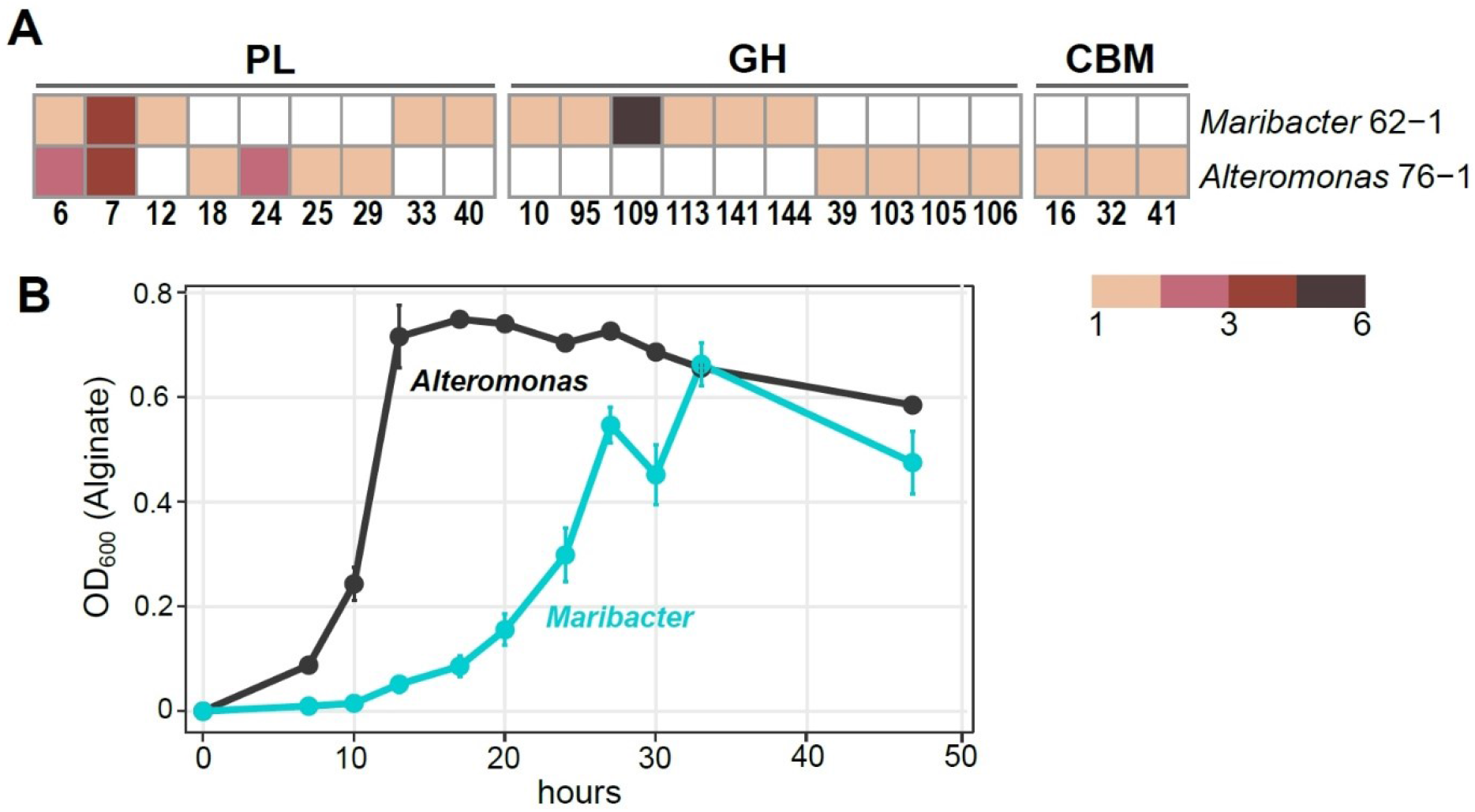
Comparison of *Maribacter dokdonensis* 62-1 with *Alteromonas* sp. 76-1 isolated from the same seawater sample. **A:** Shared and unique families of polysaccharide lyases (PL), glycoside hydrolases (GH) and carbohydrate-binding modules (CBM). **B:** Differing growth with alginate as sole carbon source.

We hypothesize that differences in CAZymes and PUL for the same substrates mediate ecological specialization. For instance, different PL combinations might specialize for different subsets of these polysaccharides, e.g. relating to polymer length, the ratio of individual carbohydrate monomers, sulfatation or the degree of side-chain decorations. For instance, the *Alteromonas*-specific PL18 might boost alginolytic activity by acting as bifunctional lyase on both poly-G and poly-M stretches (Li et al. 2011). Furthermore, strain-specific GH genes might allow the separation into distinct “polysaccharide niches” and minimize competition. For instance, a GH109 with presumed *N*-acetylgalactosaminidase activity (Lombard et al. 2014) only occurs in *Maribacter* (Fig. 5A). Finally, ecological specialization might include “temporal niche differentiation” considering variable doubling times when degrading alginate (Fig. 5B). The delayed exponential phase in *Maribacter* suggests *K*-strategy, compared to *r*-strategy in the faster-growing *Alteromonas* (Pedler et al. 2014).

## CONCLUSIONS

The diversity of CAZymes and PUL targeting alginate, ulvan and other algal carbohydrates illustrates a marked predisposition of *Maribacter dokdonensis* 62-1 for polysaccharide degradation, exemplified by distinct growth with alginate as sole carbon source. The variety of EPS-related genes might facilitate switching between free-living and surface-associated lifestyles, potentially connected to the specific colonization of algal hosts. The overall conservation of adaptive features in *Maribacter* and the “sister genus” *Zobellia* highlights their importance as macroalgal epibionts, supported by detection of PL homologs in metagenomes from brown, green and red macroalgae on global scales. The presence of both adverse (utilization of algal substrates, evading algal defense) and advantageous traits (vitamin production, biofilm control) indicates versatile lifestyles, which potentially change depending on physicochemical conditions and algal health (Kumar et al. 2016). Different hydrolytic machineries and doubling times of strain 62-1 compared to the co-occurring *Alteromonas* sp. 76-1 might avoid competition and result in colonization of discrete niches. These insights contribute to the understanding of flavobacterial CAZymes and hydrolytic activity in context of biogeochemical cycles, niche specialization and ecological interactions in the oceans.

## ACKNOWLEDGMENTS

We thank Rolf Weinert and Martin Mierzejewski for excellent laboratory assistance. Captain, crew and colleagues of RV Polarstern expedition ANTXXVIII-5 are thanked for their professional and friendly support. Sonja Voget is gratefully acknowledged for Illumina sequencing. MW was supported by the German Research Foundation (DFG) through grant WI3888/1-2.

## Author Contributions Statement

LAW conducted genomic analyses and contributed to writing. MM performed growth experiments and HPLC. JK contributed to genomic analyses. RD performed genome sequencing. MS co-organized the expedition and experiments yielding the isolation of strain 62-1. MW designed research and wrote the paper. All authors contributed to the final version of the manuscript.

## SUPPLEMENTARY FIGURES

**Fig. S1.**
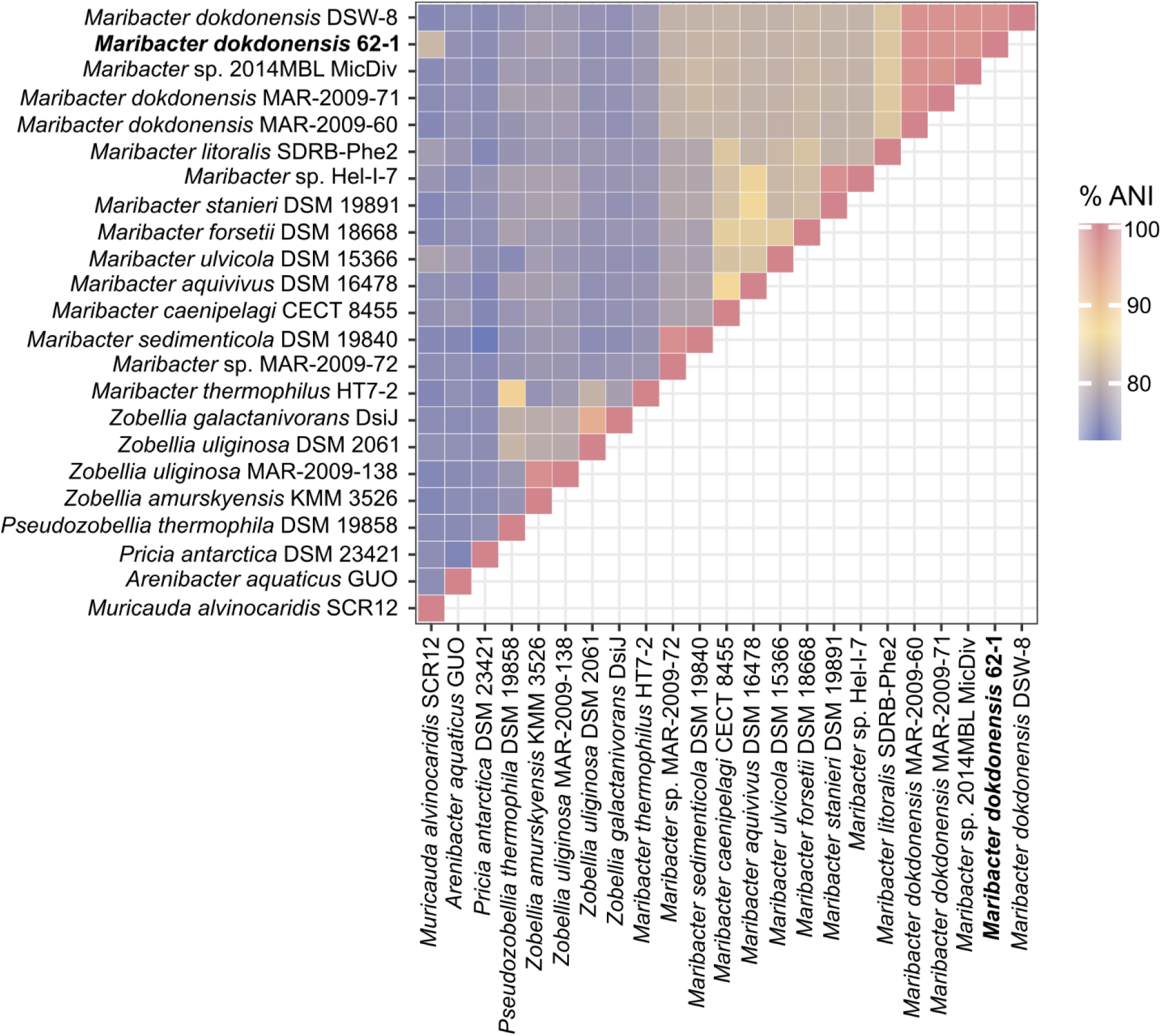
Heatmap of average nucleotide identities (ANI) between the genomes of *Maribacter dokdonensis* 62-1 and related strains.

**Fig. S2.**
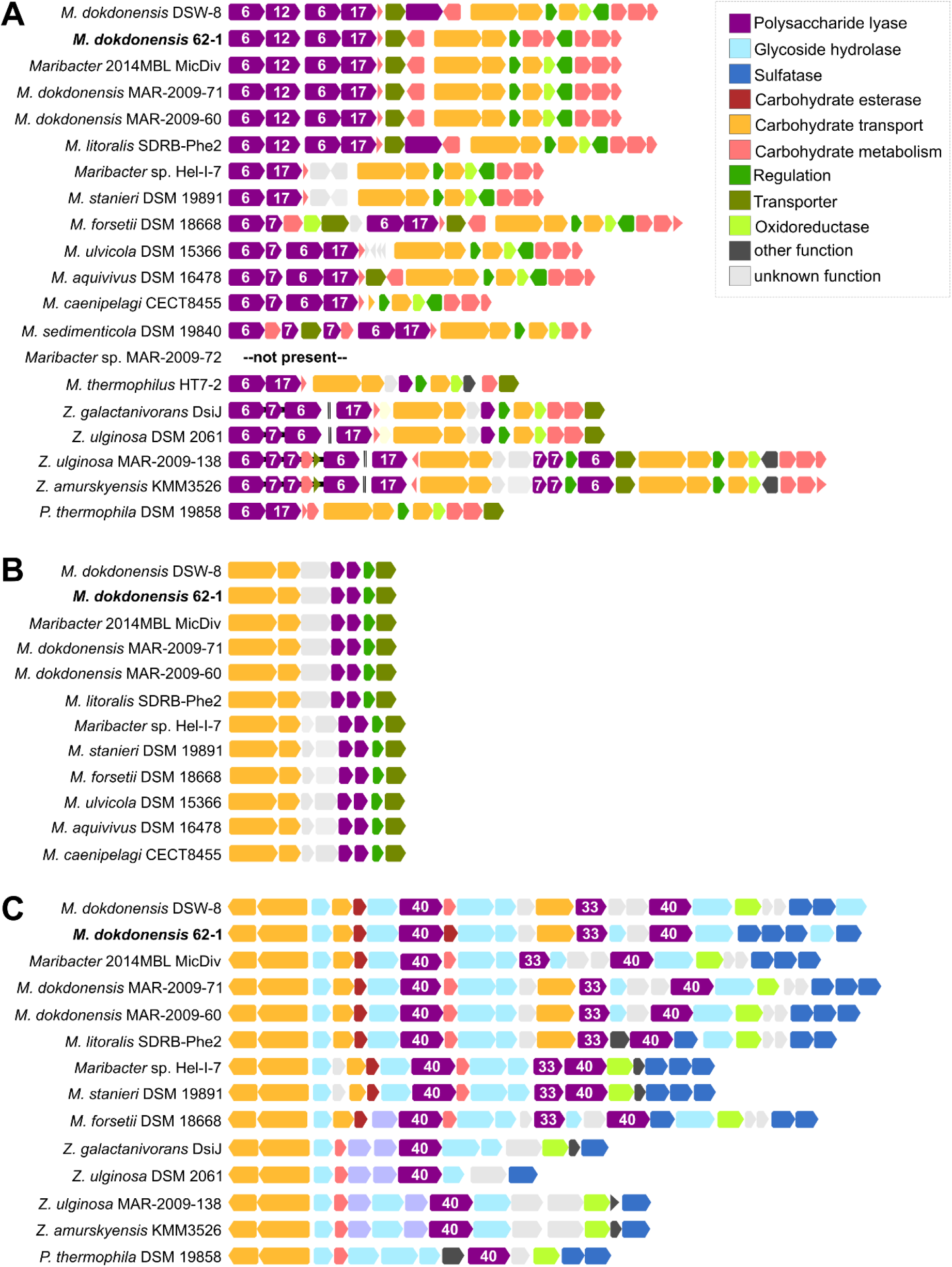
Homologs of AlgPUL1 (A), AlgPUL2 (B) and the mixed-feature PUL relating to ulvan and fucoidan (C) in genomes of *Maribacter* and *Zobellia* strains. Numbers designate PL families.

---

**Supplementary Table S1 – Strain overview.** Genome characteristics of *Maribacter dokdonensis* 62-1 and related strains used for comparative analyses.

**Supplementary Table S2 – CAZyme overview.** CAZymes, PUL, sulfatases and signal peptides in *Maribacter dokdonensis* 62-1 and related strains.

**Supplementary Table S3 – Pangenome overview.** Core, accessory and unique genes in *Maribacter dokdonensis* 62-1 and related strains.

**Supplementary Table S4 – Ecological implications.** Detection of PL homologs in microbial metagenomes from marine plants and additional algae-adaptive features of *Maribacter dokdonensis* 62-1.

